# Fungal-bacterial interaction unaffected by heatwave conditions

**DOI:** 10.64898/2026.04.29.721557

**Authors:** Maria Moreno-Druet, Sarah Pardaens, Nadejda A. Soudzilovskaia, Frederik De Laender, François Rineau

## Abstract

Climate change is reshaping soil microbial communities, yet the impact of warming in bacterial-fungal interactions (BFIs) remains underexplored. We investigated whether heatwave temperature influence BFIs and the mechanism supporting the interaction. Using co-culture experiments with two bacterial and two fungal strains isolated from heathland soil, we compared mono- and co-cultures final abundances under ambient (18°C) and heatwave (25°C) soil temperatures. Our results revealed strongly asymmetric interactions, where fungi benefited by around 5% from bacterial presence, while bacterial abundance was inhibited by around 68%, regardless of temperature. Analyses of pH confirmed that acidification by fungi was probably the main cause of this inhibition. Moreover, warming did not affect the strength or direction of these interactions, though it slightly increased fungal abundance. These findings provide direct experimental evidence that fungi can impact bacteria via acidification, and that the interaction is unaffected by temperature. Understanding these mechanisms is crucial for improving predictions of microbial community dynamics and ecosystem functioning in warming environments.

## INTRODUCTION

Climate change is altering terrestrial ecosystems at unprecedented rates with consequences for ecological processes, particularly those mediated by soil microbes, and as soils are continuously affected by environmental factors, they are especially vulnerable to accelerating impacts of climate change (Fan et al., 2024; Xu et al., 2023). Since pre-industrial times, global temperatures have risen by more than 1°C and this trend is expected to continue, resulting in more frequent and intense droughts, heatwaves, and shifts in precipitation patterns (IPCC, 2023). These environmental shifts have a direct effect on the activity and diversity of soil microbial communities, affecting crucial soil processes (De Vries & Griffiths, 2018; Ratzke et al., 2020). These communities regulate the decomposition of organic matter, drive nitrogen and phosphorus cycles, carbon sequestration, and contribute significantly to soil structure (De Vries & Griffiths, 2018; Haq et al., 2014; Nannipieri et al., 2003). Notably, estimates suggest that 80-90% of soil processes are microbially mediated (Nannipieri et al., 2003). These contributions are not simply additive, but come from the complex organization and interactions within microbial communities.

The impact soil microbes have on soil functioning is determined by their diversity and community composition (i.e. Ondik et al., 2023; Wagg et al., 2021), which is strongly driven by abiotic environmental factors. Factors such as temperature and moisture directly influence microbial growth and activity or act indirectly by altering plant inputs and soil chemistry (De Vries & Griffiths, 2018). For example, experimental soil warming often results in reduced microbial diversity and altered community composition (Y. Zhou et al., 2021), while drought stress alters both bacterial and fungal communities (Seaton et al., 2022). Among these abiotic factors, soil pH is particularly influential, as it is one of the main of factors controlling microbial metabolism and communities across ecosystems. Moreover, soil pH has consistently been identified as one of the strongest predictors of bacterial community composition, with peak diversity often found at near-neutral conditions (Fierer, 2017; Lauber et al., 2009). Nutrient availability also plays a key role: increased resource inputs, such as carbon or nitrogen pulses, typically stimulate microbial activity and can reshape community composition over time (Ratzke et al., 2020). These community-level shifts are often associated with changes in microbial biomass, taxonomic composition, and metabolic activity, which in turn alter the abundance and dominance of specific taxa (Bárcenas-Moreno et al., 2009; Seaton et al., 2022; Y. Zhou et al., 2021). In some cases, these shifts result in community simplification, functional reorganization, or dominance by stress-tolerant groups (Singh et al., 2010; J. Zhou et al., 2012; Zosso et al., 2021). Therefore, abiotic factors act as selective environmental filters, shaping microbial communities and thereby regulating soil biogeochemical processes.

Microbes rarely live in isolation; they compete for resources, prey on each other, or engage in mutualistic relations. All these types of interactions strongly influence which species eventually can coexist, thereby determining the overall composition and function of the community (Ratzke et al., 2020; Singh et al., 2010). Shifts in these microbial interactions can have cascading effects on ecosystem functioning (Y. Zhou et al., 2021). Yet despite their importance, microbial interactions have received less attention in climate change studies compared to abiotic factors (Y. Zhou et al., 2021). The effects of heat or drought on soil communities are often attributed to abiotic stress directly affecting microbial biomass, even though altered competitive or facilitative relationships among microbes, caused by the stresses, may in fact be indirectly responsible for the observed community changes or functional outcomes. Emerging evidence supports this; for instance, extreme drought was found to destabilize bacterial co-occurrence networks while leaving fungal networks relatively intact (De Vries et al., 2018). To fully understand how environmental shifts influence soil microbial communities, and thus also soil function, it is essential to take both abiotic conditions as well as interactions between microbial species into account (Singh et al., 2010).

Soil is dominated by bacteria and fungi, and their interactions, known as bacterial-fungal interactions (BFIs) are functionally relevant. These groups fulfill complementary roles: fungi play a key role in decomposing complex organic matter such as lignin and cellulose and contribute to soil structure through their hyphal networks, while bacteria are primarily involved in the decomposition of more labile substrates and drive rapid nutrient cycling and enzymatic turnover. (Haq et al., 2014; Paul, 2015). Their interactions range from competition to cooperation, including mutualism, amensalism, and neutralism (Deveau et al., 2018; Faust & Raes, 2012; Frey-Klett et al., 2011), with competition being one of the major forces shaping microbial communities (Bahram et al., 2018). In general, fungi tend to outcompete bacteria for complex substrates like cellulose, while bacteria are more competitive for simpler substrates such as glucose (Meidute et al., 2008; Wang & Kuzyakov, 2024). Mechanistically, BFI involve direct molecular interactions such as chemotaxis, antibiosis, metabolic exchange, or even the use of fungal hyphae by bacteria as dispersal vectors (Kohlmeier et al., 2005; Rousk et al., 2010). The outcomes of these interactions extend beyond microbial fitness, as they can also alter soil nutrient dynamics, resource availability, community assembly, and ecosystem functions (Deveau et al., 2018).

Despite their high ecological importance, BFIs remain underexplored. Traditional divisions between bacteriology and mycology have led to the study of bacteria and fungi as separate communities (Seaton et al., 2022), often resulting in overlooking their coexistence and complex interactions in natural environments (Carr et al., 2019; Frey-Klett et al., 2011). While recent molecular network analyses inferred microbial interactions (Yuan et al., 2021; Y. Zhou et al., 2021), the underlying mechanisms behind these interactions remain largely unexplored, especially with respect to species co-occurring in natural ecosystems. This knowledge gap constrains our ability to anticipate how microbial communities and the functions they support will respond to global change. Here, we tested the effect of increased soil temperature observed during a heatwave on the interactions between two fungal and two bacterial strains. We used in-vitro assays to measure the reciprocal effects of each bacterial species and the two fungal species at ambient and elevated temperature conditions, and examined the mechanism behind bacterial abundance reductions in cocultures. We specifically assessed the role of fungal acidification in altering substrate pH and its impact on bacterial abundance, as pH changes are a known factor influencing microbes (Fierer, 2017; Rousk et al., 2010). The study was conducted on fungal and bacteria strains naturally co-occur in European heathland ecosystems: *Umbelopsis* sp. *Cladosporium* sp., *Viridibacillus* sp., and *Bacillus mycoides*.

## MATERIAL & METHODS

### Research questions and experiment overview

To investigate how heatwave temperatures influence BFIs, we conducted controlled *in vitro* experiments comparing the final abundances of two bacterial and two fungal strains isolated from heathland soil in mono- and co-culture under two temperature conditions: 18°C (reference) and 25°C (simulated heatwave). These values correspond to soil temperatures measured at 10 cm depth in heathland soil macrocosms under a simulated heatwave in an ecotron, reflecting realistic summer temperatures under normal and elevated temperatures. We aimed to address two main research questions. **Question 1**: Does warming influence bacterial-fungal interactions (BFIs)? We hypothesized that elevated temperatures might shift interactions towards bacterial dominance due to their faster growth rates and reduced sensitivity to heatwave conditions. **Question 2**: What is the mechanism supporting the interaction ? We suspected fungal acidification of the medium to play a role. Test this, we measured growth medium pH at the end of the experiment. Additionally, we grew bacteria as monocultures at distinct pH levels to determine whether pH was the primary driver of the reduction of bacterial final abundance.

### Site description

Soil samples were collected from Mechelse Heide, a heathland ecosystem dominated by *Calluna vulgaris, Deschampsia flexuosa*, and *Molinia caerulea*, situated in National Park Hoge Kempen, Eastern Belgium. The climate of the region is characterized by a mean annual temperature of 10.3°C and 839 mm of annual precipitation. Sampling was conducted in August 29, 2023, with a 1.4 cm diameter soil core at a depth of 10 cm. A total of 30 samples were collected in a regular 5 by 4 grid pattern covering the sampling area (2 m intervals), pooled, homogenized, sieved (1.5 mm mesh), and stored at 5°C.

### Bacterial strain isolation and identification

Bacterial strains were isolated from the soil samples using the dilution plate method, where 10 g of soil was dissolved in 100 mL of phosphate buffered saline (PBS) to create a soil stock solution. A series of four tenfold dilutions (1:10, 1:100, 1:1000, and 1:10000) were made, and 200 μL from each dilution was spread onto Luria-Bertani (LB) agar for a total of 40 plates (10 per dilution). Plates were then incubated overnight at 35°C. The following day, colonies were selected based on their morphology and collected with a sterile toothpick, placed on liquid LB in sterilized glass tubes and incubated at 35°C for several days.

DNA extraction was performed on 35 randomly selected monocultures, each from different dilutions and based on different morphological characteristics, using the Norgen’s Soil DNA Isolation Plus Kit® (Norgen Biotek, Ontario, Canada), according to the manufacturer’s protocol. DNA quality was assessed using Nanodrop® analysis, and 16 bacterial isolates were selected for further analysis based on DNA quality.

PCR amplification was performed in 25 μl containing 5x Q5 reaction buffer (New England Biolabs, Ipswich, MA, USA), 10 nM dNTPs, 10 μM of both the forward (515f, 5’–GTG CCA GCM GCC GCG GTA A–3’) and reverse primer (806r, 5’–GGA CTA CHV GGG TWT CTA AT–3’), 0.02 U/μL Q5 Hot-Start High Fidelity DNA Polymerase, nuclease free water and 1 μL of DNA template using a PCR thermocycler Biorad C1000 (Biorad, Temse, Belgium). PCR conditions consisted of an initial denaturation at 98°C for 3 minutes, followed by 30 cycles of 10 seconds at 98°C, 30 seconds at 57°C, 30 seconds at 72°C and a final extension for 7 minutes at 72°C. An agarose gel electrophoresis was conducted to confirm successful PCR amplification. The PCR products (n = 16) were then sent for Sanger sequencing at Macrogen Europe (Amsterdam, the Netherlands) for identification.

For long-term storage, bacterial isolates were preserved in 50% glycerol at -80°.

### Bacterial strain selection & cultivation

We reactivated the bacterial strains *Bacillus mycoides* and *Viridibacillus* sp. from - 80°C glycerol stock, and we selected them because they are common in heathland soils, and represent functionally important bacteria. Strains were streaked onto Luria-Bertani (LB) agar plates and incubated at 35°C for 2 days. Actively growing bacterial colonies were transferred to 15 mL liquid LB medium and incubated overnight at 30°C while shaking. Optical density (OD) measurements at 600 nm were taken to confirm that the bacterial cultures were in the exponential growth phase. Following overnight growth, 1 mL of bacterial inoculum was transferred into 9 mL Modified Minimum-Melin-Norkrans (MMN) medium in sterile 15 mL falcon tubes, vortexed for 5 seconds, and incubated under shaking conditions at 30°C for 36-50 hours. This ensured that the bacteria culture reached exponential growth phase before exposure to experimental treatments. The medium composition was based on Modified Melin-Norkrans as described in Marx (1969), with nutrient adjustments, adapted to a liquid form by omitting agar, and adjusted to a pH of 6.5: KH2PO4 (0.05%), NH4Cl (0.02%), MgSO4·7H2O (0.0015%), CaCl2·2H2O (0.005%), NaCl (0.0025%), FeCl3·6H2O (0.0012%), thiamine HCl (0.0001%), and glucose (0.25%). We used MMN medium with a pH of 6.5, which is commonly used to grow fungi, but it can also support bacterial growth (Fig. S1). Its relatively low nutrient content helps to mimic soil conditions, allowing us to maintain comparable growth conditions for both fungi and bacteria and test their interactions under controlled nutrient conditions.

### Fungal strain selection

*Umbelopsis* sp. and *Cladosporium* sp. were previously were isolated from heathland soil samples in a study by Reyns (2020), and were stored at 5°C in the laboratory inventory until use. We selected these fungi based on their prevalence and relevance in heathland soils, their saprotrophic nature, and their reliable growth under controlled laboratory conditions. Both strains are commonly found in soil ecosystems, where they play key roles in decomposing organic matter.

### Fungal strain cultivation

*Umbelopsis* sp. and *Cladosporium* sp. strains were retrieved from storage at 5°C and inoculated onto 25 mL of MMN agar plates. The thicker agar (25 mL) was necessary to ensure that fungal mycelium did not submerge during the transfer to liquid media. Plates were incubated at 20°C for 8 days to ensure sufficient mycelium growth. A 7mm agar-mycelium plug was removed using a cork borer and transferred to 15 mL liquid MMN medium. Both fungal strains were given a 4-day growth advantage at 20°C to ensure enough biomass. After 4 days, the media was refreshed by carefully removing the old media using a sterile 10 mL pipette. Consistency was maintained by removing and replacing approximately 12 mL of the old medium with 12 mL of fresh medium for each sample.

### Monoculture assays

Both fungi and bacteria were cultured separately under two temperature conditions (18°C and 25°C) for 7 days to establish baseline growth data. We defined final abundance as the biomass or cell density measured at the end of the incubation period and it was quantified differently in fungi and bacteria. Bacterial final abundance was quantified by measuring OD600 using a spectrophotometer (Amersham Novaspec Plus Visible spectrophotometer). Fungal final abundance was quantified by collecting visible mycelium from the petri plates, transferring it into 15 ml falcon tubes and freezing them at -80°C overnight. Subsequently, falcon tubes were uncapped and covered with parafilm, punctured with a hole to allow for sublimation and lyophilized for 24-36 hours to remove leftover moisture and media. Following lyophilization, dry weight was measured. Optical density measurements were also collected for monoculture fungal experiments at 600 nm. This was done to correct for the contribution of particles of fungal origin to the OD. The pH of the medium was measured using a calibrated pH meter electrode at the end of the incubation period to assess any changes due to microbial activity. Initially, all culture media were adjusted to a pH of 6.5.

### Coculture assays

Coculture experiments were conducted using all possible combinations of the selected fungal and bacterial strains. Each combination was subjected to two different temperature conditions. The microbial strains used in coculture included *Umbelopsis* sp. or *Cladosporium* sp. paired with either *Viridibacillus* sp. or *Bacillus mycoides*.

Bacterial inocula were added to actively growing fungal cultures after a 4-day pre-growth period. One mL of bacterial inoculum was mixed into 12 mL of fresh MMN medium and pipetted into the petri dish containing fungal mycelium. The co-cultures were then incubated at the same temperatures (18°C and 25°C) for 7 days. Both bacterial and fungal final abundances were measured as described in the section ‘Monoculture assays’, and the pH of the medium was measured at the end of the incubation period.

### Monoculture of bacteria at different pH levels

To determine whether pH was the main driver of the reduction in bacterial abundance, we cultured the two bacterial strains in isolation under a range of pH conditions in a separate experiment, in medium with a control pH of 6.5 (the initial pH of the medium) and at adjusted pH levels of 2, 2.5, 3, and 3.5. These pH values were selected based on the degree of acidification observed in the medium after one week of fungal growth in MMN liquid. By comparing bacterial final abundances across these conditions, we aimed to determine whether the pH shifts in growth medium caused by fungal activity were responsible for the reductions in bacterial final abundances.

### Statistical analyses

We investigated the effects of temperature on bacterial-fungal interactions using a full factorial design with two factors: ‘temperature’ (with two levels: reference or 18°C, and 25°C or heatwave) and ‘Interacting species’ presence’ (with two levels: presence or absence of another strain). Additionally, we carried out an experiment with monocultures of bacteria at different pH levels to examine bacterial final abundances under varying pH at the two temperature levels but independent of ‘Interacting species presence’ treatment.

Due to the limited number of bacterial and fungal strains (two each), we fitted separate models per strain to answer question 1. In each model, we used final abundance as the response variable and included ‘temperature’ and ‘presence of interacting species’ as factors, along with their interaction. For *Viridibacillus* sp., and *Bacillus mycoides*.: ART-ANOVA (Aligned Rank Transform ANOVA) was used: because residuals from linear and generalized linear models violated normality (Shapiro-Wilk p < 0.001), and variance was not constant. For *Cladosporium* sp., and *Umbelopsis* sp: we used a linear model (LM). In each model, effect size was calculated with eta_squared() which measures the proportion of variance in the dependent variable explained by an independent variable in an ANOVA. It is computed as η^2^= SS_effect_ / SS_total_, where SS_effect_ is the sum of squares for the factor of interest, and SS_total_ is the total sum of squares in the model. Model fit was evaluated using diagnostic plots.

Subsequently, to provide context for question 2, we first tested whether temperature significantly affected pH of the growth medium within each culture type (fungus and bacteria, bacteria only, fungi only) using the Wilcoxon rank-sum test. Additionally, we assessed whether microbial abundance altered pH relative to the initial medium pH of 6.5 and we used Wilcoxon tests for each culture condition. Then, to directly address question 2, we used the function predict () based on the linear model describing the experimental outcomes of abundance of bacteria monoculture at different pH levels. We tested if pH predicted bacterial abundance reduction when in coculture with fungi. We compared these predicted values against the actual observed abundance in fungal co-culture conditions using a linear regression: log(observed abundance) ∼ log(predicted abundance). Finally, we used a R2 value to assess model fit, and to determine whether predicted values significantly deviated from observations we used Welch’s t-test and tested whether the slope and intercept of the model significantly differed from the identity line (slope = 1, intercept = 0). Diagnostic plots were assessed for model assumptions. All analyses were performed using R language of statistical computing (R Core Team, 2024).

## RESULTS

Bacterial abundance was reduced by fungal presence across all temperature conditions (Fig.1). *Bacillus mycoides* exhibited a 94% reduction in abundance when cocultured with fungi, while *Viridibacillus* sp. cocultured with fungi showed a 41% decrease compared to monoculture conditions. In contrast, fungal species abundance was not negatively affected. Fungal species even slightly increased their abundance rate when grown with bacteria, at all temperatures. *Cladosporium* sp. and *Umbelopsis* sp. increased in abundance by 6% and 4%, respectively, when cocultured with bacteria compared to them growing alone. ‘Temperature’ significantly influenced final fungal abundance. However, the interaction between ‘temperature’ and ‘interacting species’ presence’ was not significant (Table 1).

**Figure 1.**
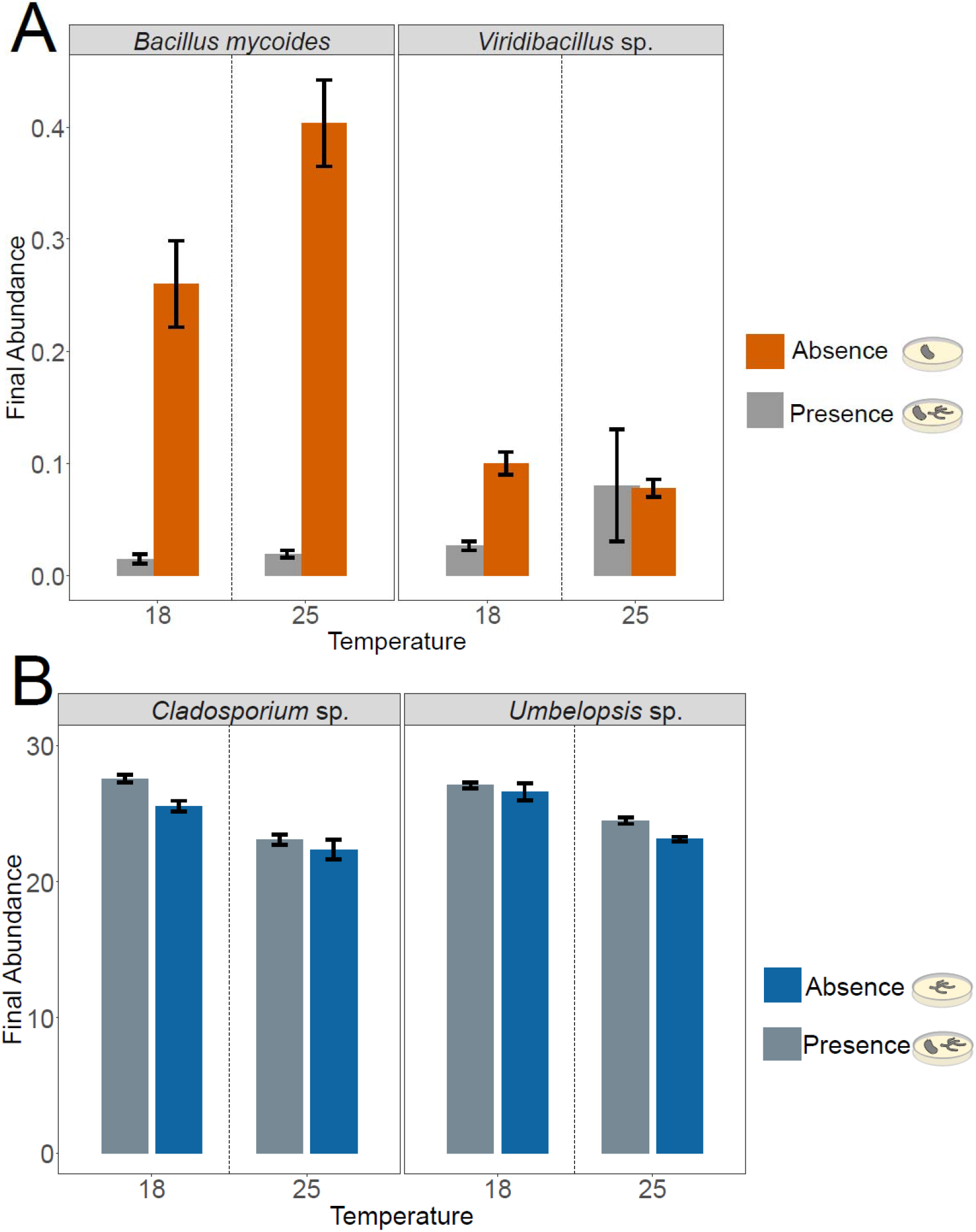
**A**. Final abundance (optical density 600 nm) of bacteria *Bacillus mycoides* and *Viridibacillus* sp. under different temperatures (18°C and 25°C) in the absence and presence of a fungal strain. Bars represent the mean final abundance (± standard error) for each bacterial strain when grown independently (absence, orange bars) and together with one fungal strain (presence, grey bars). The dotted vertical lines separate the temperature conditions. **B**. Final abundance (mg) of fungi *Cladosporium* sp. and *Umbelopsis* sp. under different temperatures (18°C and 25°C) in the absence and presence of a bacterial strain. Bars represent the mean final biomass (± standard error) for each fungal strain when grown independently (absence, blue bars) and together with a bacterial strain (presence, grey bars). The dotted vertical lines separate the temperature conditions.

**Table 1.**
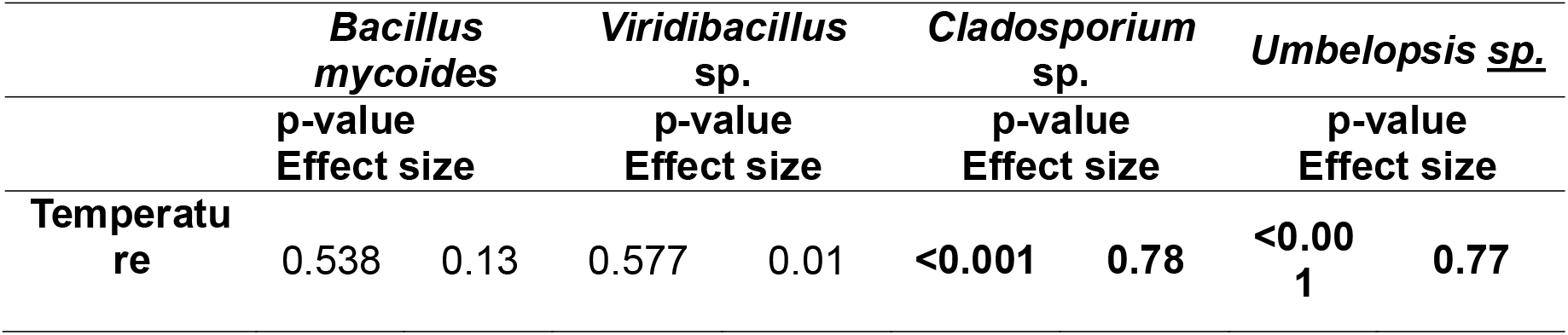
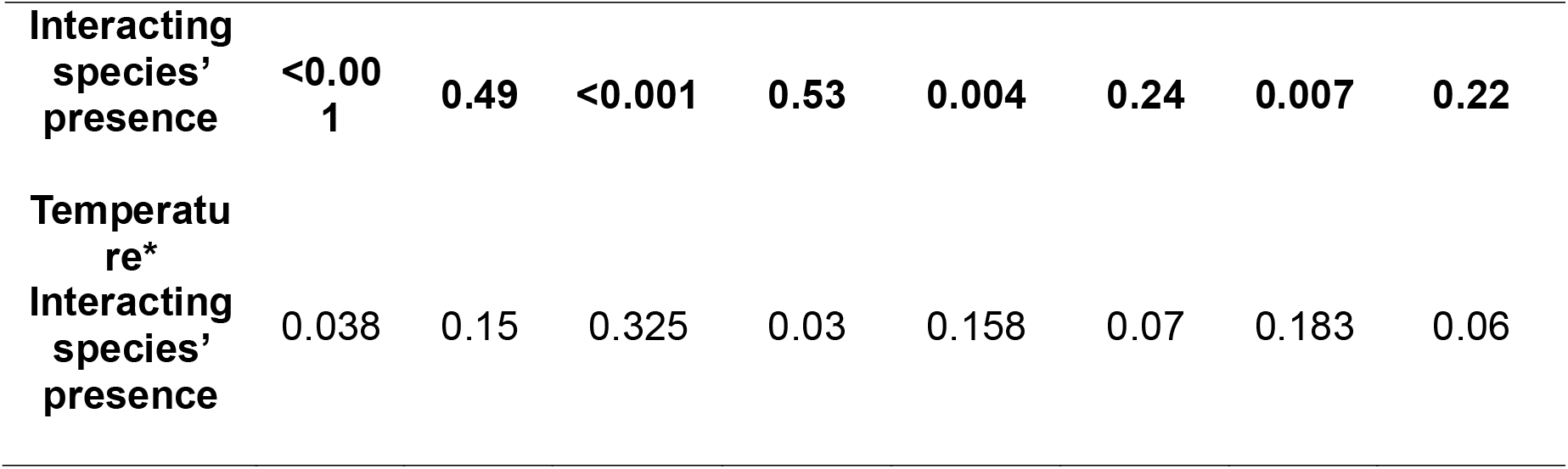
ANOVA table for each bacterial strain and the effect sizes. Significancy is **bold** (P<0.05). For each strain, we modeled final abundance as the response variable. The models included temperature, presence of interacting species, and their interaction as factors. For each strain, we used different models based on the assumptions met by the data: an ART-ANOVA for *Bacillus mycoides, Viridibacillus* sp., and a linear model (LM) for *Cladosporium* sp., and *Umbelopsis* sp. For the ART models, the effect sizes correspond to the rank-aligned transformed data. The effect sizes for the LM are on the original scale of the response variable.

Temperature significantly affected the pH of the medium only in the coculture condition, with lower values at 18°C than at 25°C (Fig.2). In contrast, no significant temperature effects were detected in fungal or bacterial monocultures (Fig.2).

**Figure 2.**
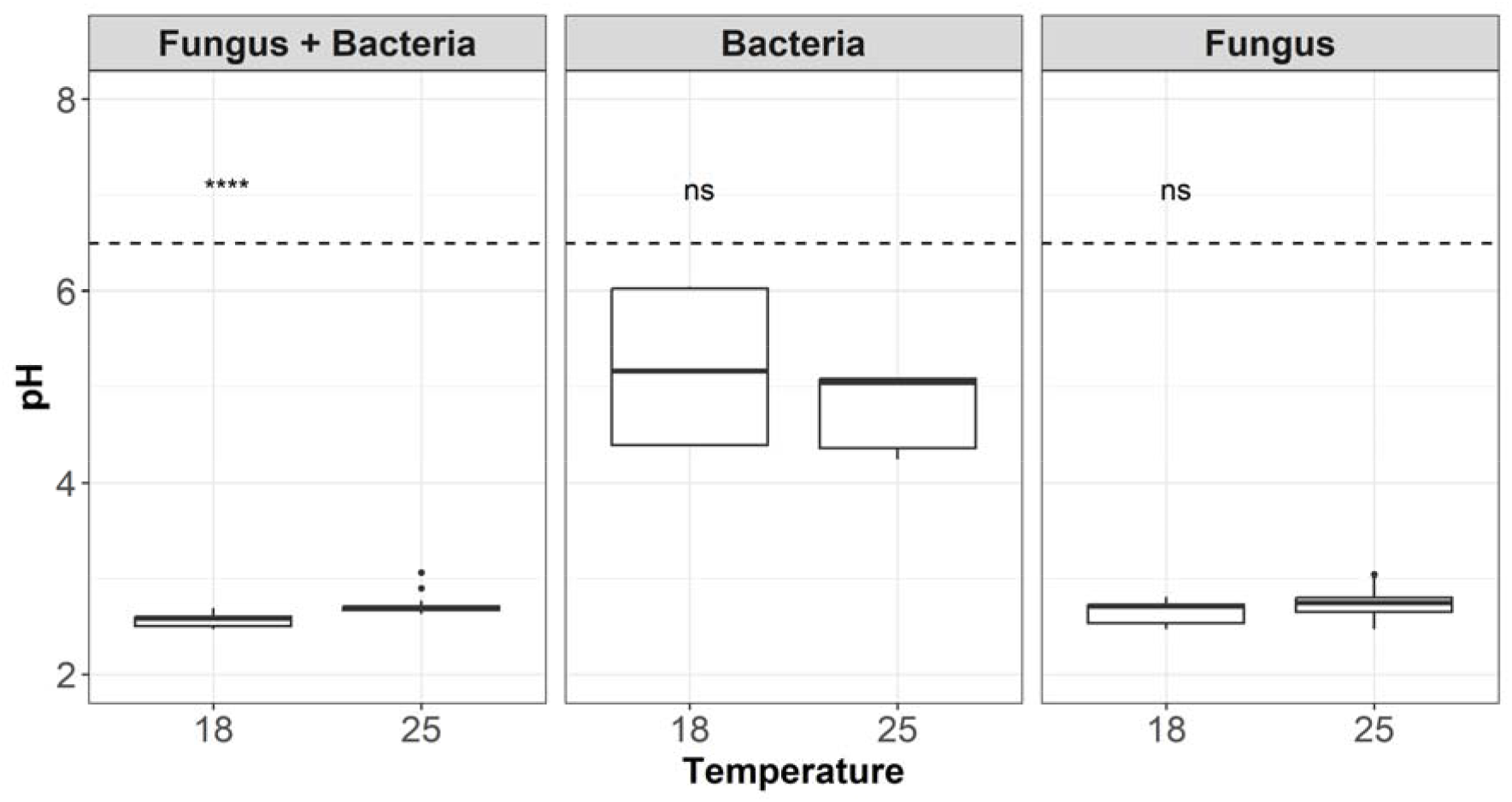
Effect of temperature on medium pH per culture type. Boxplots show pH values at two temperature conditions (18°C and 25°C) for three different setups: pH-fungus-and-bacteria, pH -bacteria (absence of fungi), and pH-fungi (absence of bacteria). A dashed horizontal line at pH = 6.5 indicates the initial pH of the medium. Statistical significance of temperature effects within each culture type (each panel) was assessed using the Wilcoxon rank-sum test, with significant differences marked by asterisks (“****” indicates a highly significant difference), while “ns” denotes no significant difference.

In all treatments, pH values of the growth medium at the end of the experiment were significantly lower than the initial medium pH (all p < 0.01). The strongest acidification occurred in treatments involving fungi, either in monoculture or in coculture. Bacterial monocultures also reduced the pH significantly, although the magnitude of change was smaller.

Bacterial final abundance was reduced due to acidification across all temperatures (Fig. 3). Optical density (OD 600) measurements showed consistent trends across pH levels (Fig. 3). Overall, a significant positive relationship was observed between bacterial optical density and initial pH levels for both *Bacillus mycoides* (R^2^ = 0.7 at 18°C, R^2^ = 0.86 at 25°C) and *Viridibacillus* sp. (R^2^ = 0.75 at 18°C, R^2^ = 0.8 at 25°C), indicating that bacterial abundance was higher in less acidic conditions.

**Figure 3.**
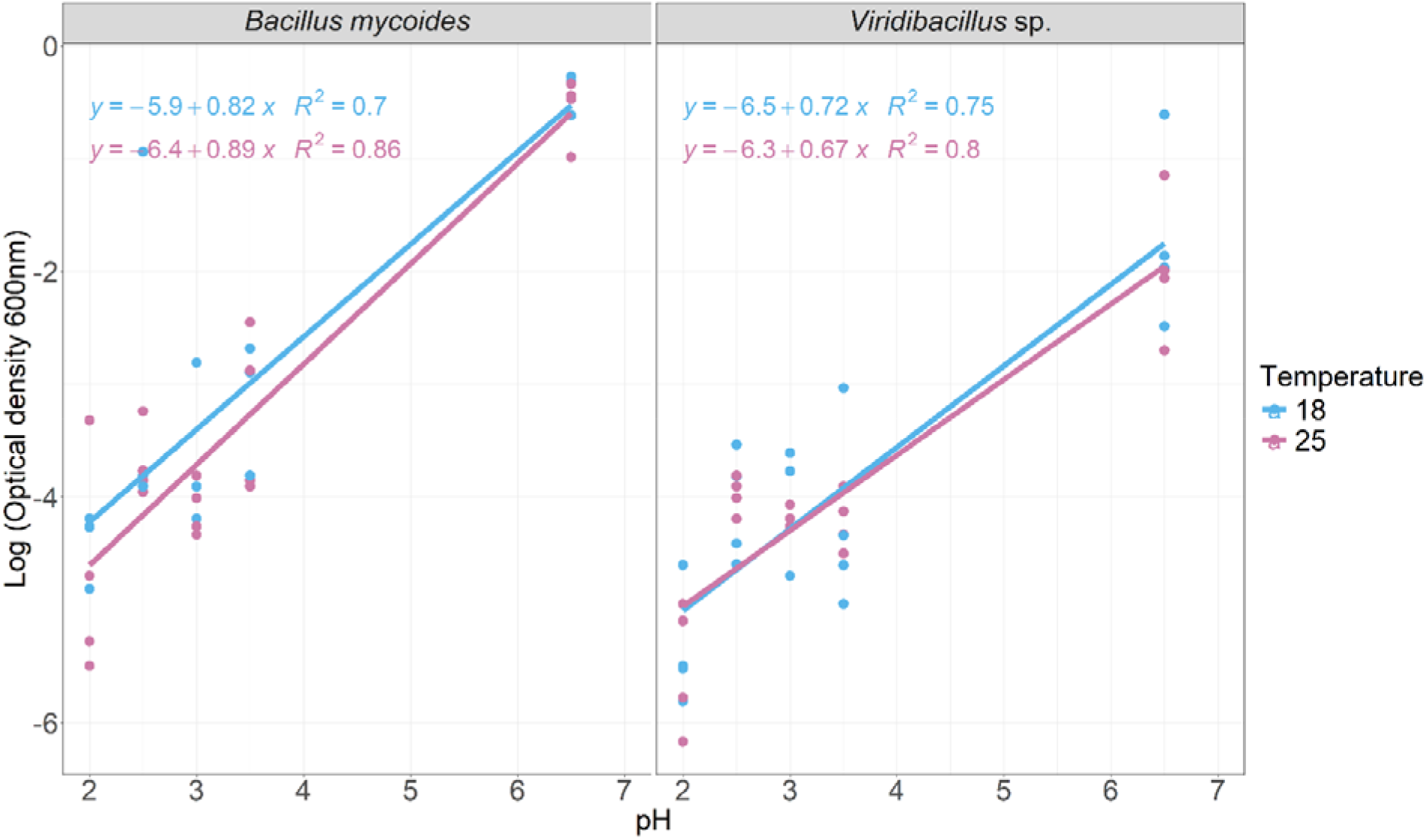
Log-transformed optical density at 600 nm (OD600) of *Viridibacillus* sp. and *Bacillus mycoides* in monocultures across a pH gradient under two temperature treatments. Linear regression lines and corresponding equations with R2 values are shown. One data point at pH 6.5 was excluded due to experimental error.

Additionally, the assessment of observed vs. predicted bacterial abundance confirmed that pH is a significant driver of bacterial final abundance (Fig.4); for *Bacillus mycoides* regression analysis of log-transformed predicted and observed abundances revealed a strong positive relationship (R^2^ = 0.66, p < 0.001). While the slope was significantly different from zero (p < 0.001), it did not differ significantly from 1 (p = 0.20), and the intercept did not differ significantly from 0 (p = 0.96) (Fig.4). However, a joint hypothesis (testing together slope and intercept) test showed that the overall regression line differed significantly from the 1:1 identity line (p = 0.0014). For *Viridibacillus sp*., predicted and observed log-transformed abundances had a significant positive relationship (R^2^ = 0.36, p < 0.001) (Fig.4). In the fitted regression, both the slope and the intercept significantly different from the 1:1 line (slope ≠ 1: p < 0.001; intercept ≠ 0: p 0.001). Additionally, a joint hypothesis test confirmed that the regression line significantly deviated from the 1:1 line (p < 0.001), indicating an overprediction by the pH-based model (Fig. 4).

**Figure 4.**
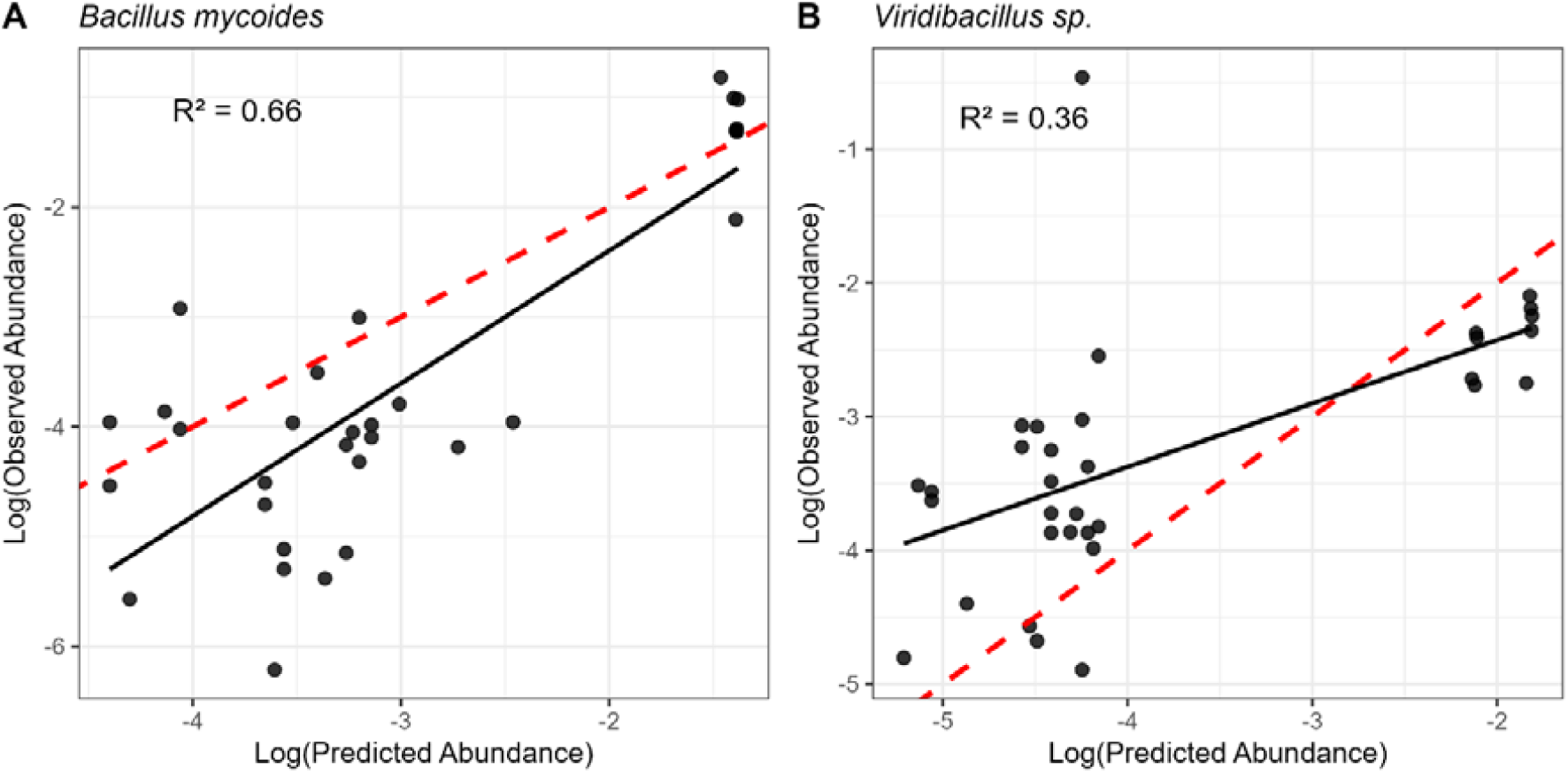
Observed abundance vs. the abundance predicted from pH. The black line represents the linear regression model fit; the red dashed line indicates the identity line (slope = 1, intercept = 0). Each point represents an individual observation. For *Bacillus mycoides*, the regression slope did not differ significantly from 1, whereas for *Viridibacillus* sp. the slope was significantly different from 1. One data point at pH 6.5 was excluded due to experimental error.

## DISCUSSION

Microbial interactions play a crucial role in ecosystems stability and will determine the composition and functioning of microbial communities, ultimately influencing nutrient cycling (Osburn et al., 2023; Yuan et al., 2021). While numerous studies have examined the effects of climate change on microbial communities, much less is known about its influence on microbial interactions. In particular, it remains largely unexplored how climate change alters these interactions, as most existing insights come from molecular network inferences rather than direct measurements. Our study is among the first to address this knowledge gap for microbial species co-existing in a natural ecosystem. In cocultures, we observed that fungi benefited from bacteria, with a modest gain of about 5%, whereas bacterial abundance was strongly inhibited by approximately 68%. This implies that the interaction type we detected is closer to amensalism (0/-) than to parasitism and/or predation (-/+), and that this phenomenon occurred independently of temperature conditions.

### Fungi suppress bacteria with minimal gain: evidence of amensalism

Amensalism describes an interaction where one species is harmed while the other is unaffected. Fungi can impact bacteria by rapidly colonizing available niches and depleting resources limiting bacterial abundance. In fact, some fungi produce hydrolytic enzymes like proteases and chitinases to break down proteins and effectively inhibit other organisms’ growth (Wang & Kuzyakov, 2024). They can also produce antimicrobial substances such as antibiotics, acids and secondary metabolites that inhibit bacterial growth (i.e. Kwak et al., 2016; Mathivanan et al., 2008; Wei et al., 2022). While these mechanisms are well-documented most studies do not measure whether the fungi benefit or remain unaffected. There are examples of amensalism in yeast-yeast or bacteria-bacteria systems and they are often related to food fermentation (García et al., 2017, 2019; Pommier et al., 2005). Therefore, direct experimental evidence of soil fungal-bacterial amensalism is scarce.

Although our findings point towards amensalism we still detected a slight increase in fungal abundance, and other studies show that fungi can also exploit bacteria more directly. For example, *Morchella crassipes*, a saprotrophic and ectomycorrhizal soil fungus has been proven to farm, and consequently, consume bacteria or bacterial exudates. In that specific scenario, bacterial and fungal species act as mutualists until medium resources are limited and, then, the fungus uses the farmed bacterium for its own benefit (Pion et al., 2013). Additionally, Basidiomycota fungi may also feed on bacterial biomass, particularly after glucose is depleted (Bastida et al., 2019). These examples represent predation or parasitism (-/+), where fungi obtain a direct nutritional benefit from bacteria. In contrast, the results observed in this study, where fungi suppressed bacterial growth while obtaining minimal or no direct benefit, supports the interpretation of amensalism.

### Temperature effect on BFI

In monocultures, temperature increase affected only fungal abundance. Additionally, in none of the cases the interaction between temperature and bacteria presence had an impact on fungal abundance. While both bacteria and fungi from temperate heathlands are mesophilic and typically grow well between 20°C and 30°C (Money, 2024; Pietikäinen et al., 2005), the observed increase in fungal abundance at 25°C suggests a stronger temperature responsiveness in fungi under our experimental conditions. Nevertheless, bacterial abundance remained stable across temperatures, and it was always affected by fungi regardless of temperature treatment which may indicate that factors other than thermal optima, such as competition with fungi or resource partitioning, limited their response.

### Role of fungal-induced substrate acidification

We found that the lowest pH values measured at the end of the experiments were observed in conditions where fungi were present, either alone or in coculture. We also found a strong positive correlation between the observed bacterial abundance in the interaction experiment and the predicted abundance of bacteria based on the bacterial monoculture-pH experiment across both temperatures and bacterial strains. This proved that the pH of the medium was an important driver in bacterial abundance reduction. For *Bacillus mycoides*, abundance patterns closely followed predictions based on pH, suggesting a strong sensitivity to acidification. However, the observed suppression in co-culture conditions exceeded what would be expected from pH alone, indicating that other factors, such as competition for nutrients, production of antagonistic compounds or fungi feeding on bacterial biomass, may be involved. In contrast, the response of *Viridibacillus* sp. was less predictable, but still significant. Its abundance showed a weaker correlation with pH, and the deviation from predicted values suggested that this response may be explained by other factors. This could include fungal-derived metabolites or altered resource availability, which are not captured by pH effects alone. Considering earlier works that reported intolerance of bacteria to acidic conditions (Fierer, 2017; Li et al., 2023; Rousk et al., 2010), our results suggests that the main, but not only, mechanism through which fungi outcompeted bacteria had to do with substrate acidification.

Specifically, fungi are known to produce organic acids as byproducts (Wang and Kuzyakov, 2024). These acids can locally lower the pH of the medium, creating an environment that may inhibit bacterial activity and growth. Oxalic acid, a low molecular weight organic acid, is among the most relevant fungal metabolite and can be metabolized by certain bacteria; it represents a key biochemical agent in shaping BFIs (Deveau et al., 2018; Palmieri et al., 2019). In fact, it seems to act as a signaling molecule for bacteria to find fungi to interact (Rudnick et al., 2015). However, organic acids produced by *Lentinula edodes*, particularly oxalic acid, exhibited significant antibacterial activity against various phytopathogenic bacteria (Kwak et al., 2016). Thus, the extent to which oxalic acid contributes to BFI, especially under heatwave conditions, remains unexplored and could provide a mechanistic explanation for the observed bacterial suppression in co-cultures.

Moreover, pH has been identified as a more critical determinant of bacterial community composition than other environmental factors (Fierer, 2017; Li et al., 2023; Rousk et al., 2010). However, there is also evidence that bacteria can negatively affect fungi (Höppener-Ogawa et al., 2009; Liu et al., 2016; Y. Zhou et al., 2022), which we did not observe in our experiment. It is important to note that the bacterial and fungal strains used here are common heathland soil isolates, suggesting that the lack of bacterial inhibition on fungi in our system may be representative of natural conditions in heathland soils. This contrasts with studies on mycorrhizal fungi, where certain bacteria have been shown to promote fungal performance and symbiosis (Deveau & Labbé, 2016). Thus, the direction and strength of bacterial-fungal interactions may vary substantially depending on the fungal functional group and ecological context, highlighting the complexity of microbial relationships in soil.

### Implications for soil microbial structure, biochemical cycles, and ecosystem functioning

In heathland ecosystems, shifts in the fungal-bacterial proportions due to climate change can affect plant communities and nutrient cycling. Heathland soils range from acidic (<4.5) to weakly acidic (4.5-6) and species are adapted to these acidic conditions (De Graaf et al., 2009). Additionally, warming and drought have been proved to alter the microbial communities and nutrient availability in temperate heathlands (Haugwitz et al., 2014). Nevertheless, interactions have not been that widely studied and, therefore future predictions about soil microorganisms under climate change are complex, especially because while some researchers expect that under warming, microbial interactions would shift towards antagonism (as observed by Reyns in laboratory experiments specifically examining fungus-fungus interactions), others suggest that cooperative interactions would be the norm (Reyns, 2020; Wang & Kuzyakov, 2024).

Previous studies suggested that rising temperatures tend to favor bacterial competitive abilities, while acidification, particularly from anthropogenic sources such as fertilization, can favor fungi (Rousk et al., 2010; Wang & Kuzyakov, 2024). The fact that both bacterial strains were suppressed in coculture, with effects partly but not entirely explained by pH, suggests that multiple fungal-mediated mechanisms may act simultaneously. Our results evidence that fungi impact bacteria via acidification of substrate, regardless of whether a heatwave occurs.

### Limitations & future directions

We examined four bacterial strains under a single environmental factor (temperature) with two levels and with a single competitor in each case, using petri dishes. These conditions do not represent natural environments where interactions are more complex but these simplified experiments help us explain nature and disentangle smaller scale mechanisms (Drake & Kramer, 2012). Microbial interactions can be highly context-dependent, changing with environmental conditions, community composition, or the medium in which they take place. Thus, future studies should explore more strains, more realistic conditions such as soil microcosms, and a wider range of environmental factors.

## Conclusion

We examined whether heatwaves modified bacterial-fungal interactions and we addressed the limitations of network-based molecular studies (as they often struggle to distinguish between microbial interactions from habitat selection effects). Moreover, by applying direct co-cultivation experiments, we provided novel insights into microbial interactions in controlled environments. In particular, we showed direct evidence of asymmetric fungal-bacterial interactions isolated from a heathland soil, where fungi inhibited bacterial growth while benefiting slightly from bacterial presence, possible due to fungal acidification of the medium. These findings highlight the complexity of microbial dynamics in soil environments, suggesting that both abiotic factors and microbial interactions must be considered together to improve predictions of ecosystem responses to environmental changes.

## Acknowledgements

This study was financially supported by BOF (Bijzonder Onderzoeksfonds) and FSR (Fonds Spécial de Recherche) (both Special Research Funds) from Hasselt University and Namur University, respectively, under the research grant number BOF21DOCNA01

---

**Figure S1.**
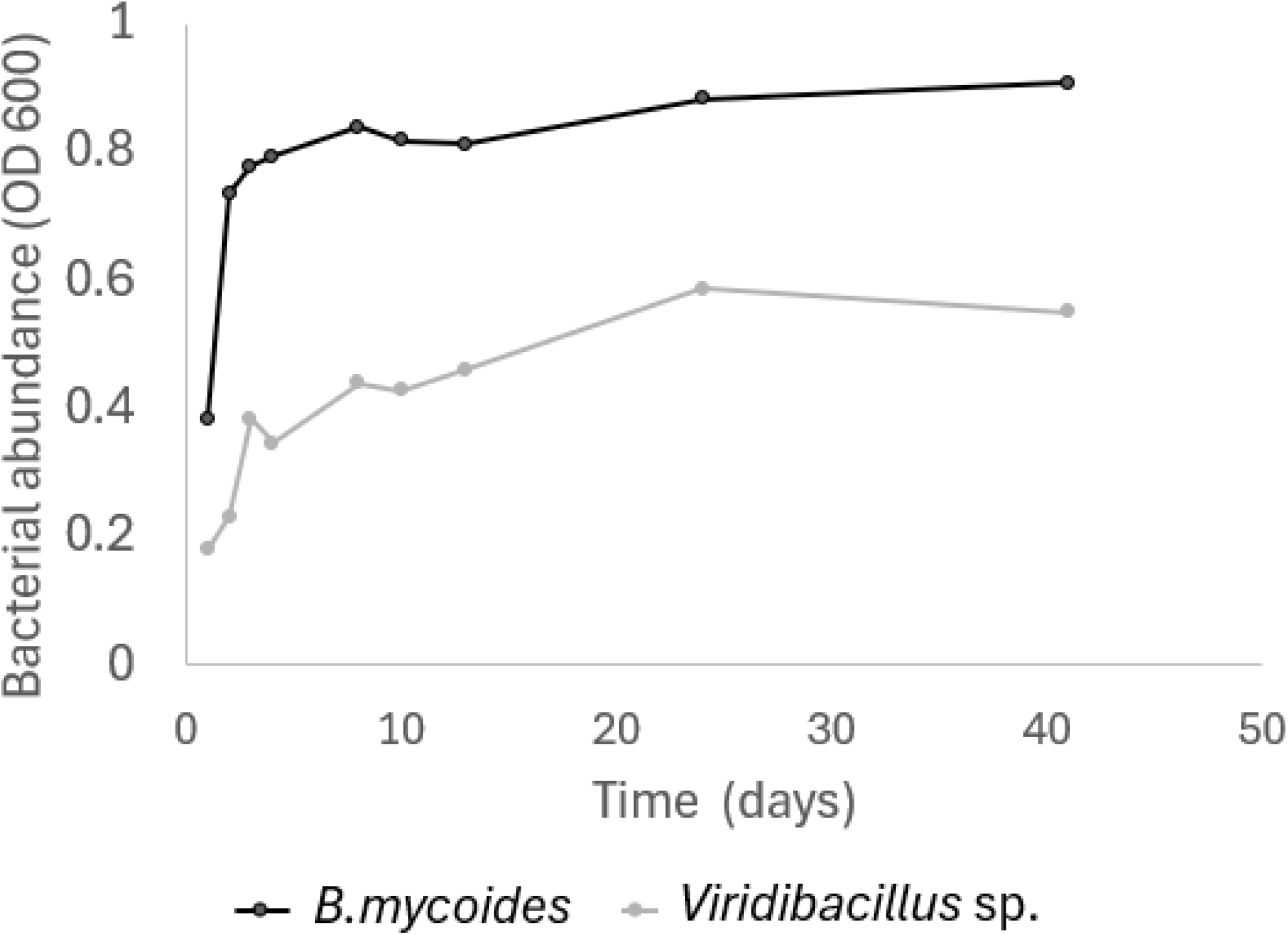
Growth dynamics of *B*.*mycoides* and *Viridibacillus*.

## Notes

**Conflict of interest** The authors declare no conflicts of interest.

### Competing Interest Statement

The authors have declared no competing interest.

## REFERENCES

Bahram, M., Hildebrand, F., Forslund, S. K., Anderson, J. L., Soudzilovskaia, N. A., Bodegom, P. M., Bengtsson-Palme, J., Anslan, S., Coelho, L. P., Harend, H., Huerta-Cepas, J., Medema, M. H., Maltz, M. R., Mundra, S., Olsson, P. A., Pent, M., Põlme, S., Sunagawa, S., Ryberg, M., … Bork, P. (2018). Structure and function of the global topsoil microbiome. Nature, 560(7717), 233–237. 10.1038/s41586-018-0386-6

Bárcenas-Moreno, G., Gómez-Brandón, M., Rousk, J., & Bååth, E. (2009). Adaptation of soil microbial communities to temperature: Comparison of fungi and bacteria in a laboratory experiment. Global Change Biology, 15(12), 2950–2957. 10.1111/j.1365-2486.2009.01882.x

Bastida, F., García, C., Fierer, N., Eldridge, D. J., Bowker, M. A., Abades, S., Alfaro, F. D., Asefaw Berhe, A., Cutler, N. A., Gallardo, A., García-Velázquez, L., Hart, S. C., Hayes, P. E., Hernández, T., Hseu, Z.-Y., Jehmlich, N., Kirchmair, M., Lambers, H., Neuhauser, S., … Delgado-Baquerizo, M. (2019). Global ecological predictors of the soil priming effect. Nature Communications, 10(1), 3481. 10.1038/s41467-019-11472-7

Carr, A., Diener, C., Baliga, N. S., & Gibbons, S. M. (2019). Use and abuse of correlation analyses in microbial ecology. The ISME Journal, 13(11), 2647– 2655. 10.1038/s41396-019-0459-z

De Graaf, M. C. C., Bobbink, R., Smits, N. A. C., Van Diggelen, R., & Roelofs, Jan. G.M. (2009). Biodiversity, vegetation gradients and key biogeochemical processes in the heathland landscape. Biological Conservation, 142(10), 2191–2201. 10.1016/j.biocon.2009.04.020

De Vries, F. T., & Griffiths, R. I. (2018). Impacts of Climate Change on Soil Microbial Communities and Their Functioning. In Developments in Soil Science (Vol. 35, pp. 111–129). Elsevier. 10.1016/B978-0-444-63865-6.00005-3

De Vries, F. T., Griffiths, R. I., Bailey, M., Craig, H., Girlanda, M., Gweon, H. S., Hallin, S., Kaisermann, A., Keith, A. M., Kretzschmar, M., Lemanceau, P., Lumini, E., Mason, K. E., Oliver, A., Ostle, N., Prosser, J. I., Thion, C., Thomson, B., & Bardgett, R. D. (2018). Soil bacterial networks are less stable under drought than fungal networks. Nature Communications, 9(1), 3033. 10.1038/s41467-018-05516-7

Deveau, A., Bonito, G., Uehling, J., Paoletti, M., Becker, M., Bindschedler, S., Hacquard, S., Hervé, V., Labbé, J., Lastovetsky, O. A., Mieszkin, S., Millet, L. J., Vajna, B., Junier, P., Bonfante, P., Krom, B. P., Olsson, S., Van Elsas, J. D., & Wick, L. Y. (2018). Bacterial–fungal interactions: Ecology, mechanisms and challenges. FEMS Microbiology Reviews, 42(3), 335–352. 10.1093/femsre/fuy008

Deveau, A., & Labbé, J. (2016). Mycorrhiza helper bacteria. In F. Martin (Ed.), Molecular Mycorrhizal Symbiosis (1st edn, pp. 437–450). Wiley. 10.1002/9781118951446.ch24

Drake, J. M., & Kramer, A. M. (2012). Mechanistic analogy: How microcosms explain nature. Theoretical Ecology, 5(3), 433–444. 10.1007/s12080-011-0134-0

Fan, X., Zhang, Y., Shi, K., Peng, J., Liu, Y., Zhou, Y., Liu, Y., Zhu, Q., Song, C., Wan, R., Zhao, X., & Woolway, R. I. (2024). Surging compound drought–heatwaves underrated in global soils. 121(42).

Faust, K., & Raes, J. (2012). Microbial interactions: From networks to models. Nature Reviews Microbiology, 10(8), 538–550. 10.1038/nrmicro2832

Fierer, N. (2017). Embracing the unknown: Disentangling the complexities of the soil microbiome. Nature Reviews Microbiology, 15(10), 579–590. 10.1038/nrmicro.2017.87

Frey-Klett, P., Burlinson, P., Deveau, A., Barret, M., Tarkka, M., & Sarniguet, A. (2011). Bacterial-Fungal Interactions: Hyphens between Agricultural, Clinical, Environmental, and Food Microbiologists. Microbiology and Molecular Biology Reviews, 75(4), 583–609. 10.1128/MMBR.00020-11

García, C., Rendueles, M., & Díaz, M. (2017). Microbial amensalism in Lactobacillus casei and Pseudomonas taetrolens mixed culture. Bioprocess and Biosystems Engineering, 40(7), 1111–1122. 10.1007/s00449-017-1773-3

García, C., Rendueles, M., & Díaz, M. (2019). Liquid-phase food fermentations with microbial consortia involving lactic acid bacteria: A review. Food Research International, 119, 207–220. 10.1016/j.foodres.2019.01.043

Haq, I. U., Zhang, M., Yang, P., & Van Elsas, J. D. (2014). The Interactions of Bacteria with Fungi in Soil. In Advances in Applied Microbiology (Vol. 89, pp. 185–215). Elsevier. 10.1016/B978-0-12-800259-9.00005-6

Haugwitz, M. S., Bergmark, L., Priemé, A., Christensen, S., Beier, C., & Michelsen, A. (2014). Soil microorganisms respond to five years of climate change manipulations and elevated atmospheric CO2 in a temperate heath ecosystem. Plant and Soil, 374(1–2), 211–222. 10.1007/s11104-013-1855-1

Höppener-Ogawa, S., Leveau, J. H. J., Van Veen, J. A., & De Boer, W. (2009). Mycophagous growth of Collimonas bacteria in natural soils, impact on fungal biomass turnover and interactions with mycophagous Trichoderma fungi. The ISME Journal, 3(2), 190–198. 10.1038/ismej.2008.97

IPCC. (2023). Climate Change 2023: Synthesis Report. Contribution of Working Groups I, II and III to the Sixth Assessment Report of the Intergovernmental Panel on Climate Change.

Kohlmeier, S., Smits, T. H. M., Ford, R. M., Keel, C., Harms, H., & Wick, L. Y. (2005). Taking the Fungal Highway: Mobilization of Pollutant-Degrading Bacteria by Fungi. Environmental Science & Technology, 39(12), 4640–4646. 10.1021/es047979z

Kwak, A.-M., Lee, I.-K., Lee, S.-Y., Yun, B.-S., & Kang, H.-W. (2016). Oxalic Acid from Lentinula edodes Culture Filtrate: Antimicrobial Activity on Phytopathogenic Bacteria and Qualitative and Quantitative Analyses. Mycobiology, 44(4), 338– 342. 10.5941/MYCO.2016.44.4.338

Lauber, C. L., Hamady, M., Knight, R., & Fierer, N. (2009). Pyrosequencing-Based Assessment of Soil pH as a Predictor of Soil Bacterial Community Structure at the Continental Scale. Applied and Environmental Microbiology, 75(15), 5111– 5120. 10.1128/AEM.00335-09

Li, X., Chen, D., Carrión, V. J., Revillini, D., Yin, S., Dong, Y., Zhang, T., Wang, X., & Delgado-Baquerizo, M. (2023). Acidification suppresses the natural capacity of soil microbiome to fight pathogenic Fusarium infections. Nature Communications, 14(1), 5090. 10.1038/s41467-023-40810-z

Liu, K., Hu, H., Wang, W., & Zhang, X. (2016). Genetic engineering of Pseudomonas chlororaphis GP72 for the enhanced production of 2-Hydroxyphenazine. Microbial Cell Factories, 15(1), 131. 10.1186/s12934-016-0529-0

Marx, D. H. (1969). The influence of ectotrophic mycorrhizal fungi on the resistance of pine roots to pathogenic infections. I. Antagonism of mycorrhizal fungi to root pathogenic fungi and soil bacteria. Phytopathology, 59, 153–163.

Mathivanan, N., Prabavathy, V. R., & Vijayanandraj, V. R. (2008). The Effect of Fungal Secondary Metabolites on Bacterial and Fungal Pathogens. In P. Karlovsky (Ed.), Secondary Metabolites in Soil Ecology (Vol. 14, pp. 129–140). Springer Berlin Heidelberg. 10.1007/978-3-540-74543-3_7

Meidute, S., Demoling, F., & Bååth, E. (2008). Antagonistic and synergistic effects of fungal and bacterial growth in soil after adding different carbon and nitrogen sources. Soil Biology and Biochemistry, 40(9), 2334–2343. 10.1016/j.soilbio.2008.05.011

Money, N. P. (2024). Fungal thermotolerance revisited and why climate change is unlikely to be supercharging pathogenic fungi (yet). Fungal Biology, 128(1), 1638–1641. 10.1016/j.funbio.2024.01.005

Nannipieri, P., Ascher, J., Ceccherini, M. T., Landi, L., Pietramellara, G., & Renella, G. (2003). Microbial diversity and soil functions. European Journal of Soil Science, 54(4), 655–670. 10.1046/j.1351-0754.2003.0556.x

Ondik, M. M., Ooi, M. K. J., & Muñoz-Rojas, M. (2023). Soil microbial community composition and functions are disrupted by fire and land use in a Mediterranean woodland. Science of The Total Environment, 895, 165088. 10.1016/j.scitotenv.2023.165088

Osburn, E. D., Yang, G., Rillig, M. C., & Strickland, M. S. (2023). Evaluating the role of bacterial diversity in supporting soil ecosystem functions under anthropogenic stress. ISME Communications, 3(1), 66. 10.1038/s43705-023-00273-1

Palmieri, F., Estoppey, A., House, G. L., Lohberger, A., Bindschedler, S., Chain, P. S. G., & Junier, P. (2019). Oxalic acid, a molecule at the crossroads of bacterial-fungal interactions. In Advances in Applied Microbiology (Vol. 106, pp. 49–77). Elsevier. 10.1016/bs.aambs.2018.10.001

Paul, E. A. (2015). Soil microbiology, ecology and biochemistry (4th edition). Academic Press is an imprint of Elsevier.

Pietikäinen, J., Pettersson, M., & Bååth, E. (2005). Comparison of temperature effects on soil respiration and bacterial and fungal growth rates. FEMS Microbiology Ecology, 52(1), 49–58. 10.1016/j.femsec.2004.10.002

Pion, M., Spangenberg, J. E., Simon, A., Bindschedler, S., Flury, C., Chatelain, A., Bshary, R., Job, D., & Junier, P. (2013). Bacterial farming by the fungus Morchella crassipes. Proceedings of the Royal Society B: Biological Sciences, 280(1773), 20132242. 10.1098/rspb.2013.2242

Pommier, S., Strehaiano, P., & Delia, M. (2005). Modelling the growth dynamics of interacting mixed cultures: A case of amensalism. International Journal of Food Microbiology, 100(1–3), 131–139. 10.1016/j.ijfoodmicro.2004.10.010

R Core Team. (2024). R: A Language and Environment for Statistical Computing (Version 4.4.2) [Computer software]. https://www.R-project.org/

Ratzke, C., Barrere, J., & Gore, J. (2020). Strength of species interactions determines biodiversity and stability in microbial communities. Nature Ecology & Evolution, 4(3), 376–383. 10.1038/s41559-020-1099-4

Reyns, W. (2020). Soil carbon sequestration in heathlands the effects of climate change on fungi. University of Hasselt, University of Namur.

Rousk, J., Bååth, E., Brookes, P. C., Lauber, C. L., Lozupone, C., Caporaso, J. G., Knight, R., & Fierer, N. (2010). Soil bacterial and fungal communities across a pH gradient in an arable soil. The ISME Journal, 4(10), 1340–1351. 10.1038/ismej.2010.58

Rudnick, M. B., Van Veen, J. A., & De Boer, W. (2015). Oxalic acid: A signal molecule for fungus-feedingbacteria of the genus Collimonas? Environmental Microbiology Reports, 7(5), 709–714.

Seaton, F. M., Reinsch, S., Goodall, T., White, N., Jones, D. L., Griffiths, R. I., Creer, S., Smith, A., Emmett, B. A., & Robinson, D. A. (2022). Long-Term Drought and Warming Alter Soil Bacterial and Fungal Communities in an Upland Heathland. Ecosystems, 25(6), 1279–1294. 10.1007/s10021-021-00715-8

Singh, B. K., Bardgett, R. D., Smith, P., & Reay, D. S. (2010). Microorganisms and climate change: Terrestrial feedbacks and mitigation options. Nature Reviews Microbiology, 8(11), 779–790. 10.1038/nrmicro2439

Wagg, C., Hautier, Y., Pellkofer, S., Banerjee, S., Schmid, B., & Van Der Heijden, M.G. (2021). Diversity and asynchrony in soil microbial communities stabilizes ecosystem functioning. eLife, 10, e62813. 10.7554/eLife.62813

Wang, C., & Kuzyakov, Y. (2024). Mechanisms and implications of bacterial–fungal competition for soil resources. The ISME Journal, 18(1), wrae073. 10.1093/ismejo/wrae073

Wei, L., Zhang, Q., Xie, A., Xiao, Y., Guo, K., Mu, S., Xie, Y., Li, Z., & He, T. (2022). Isolation of Bioactive Compounds, Antibacterial Activity, and Action Mechanism of Spore Powder From Aspergillus niger xj. Frontiers in Microbiology, 13, 934857. 10.3389/fmicb.2022.934857

Xu, H., Huang, L., Chen, J., Zhou, H., Wan, Y., Qu, Q., Wang, M., & Xue, S. (2023). Changes in soil microbial activity and their linkages with soil carbon under global warming. CATENA, 232, 107419. 10.1016/j.catena.2023.107419

Yuan, M. M., Guo, X., Wu, L., Zhang, Y., Xiao, N., Ning, D., Shi, Z., Zhou, X., Wu, L., Yang, Y., Tiedje, J. M., & Zhou, J. (2021). Climate warming enhances microbial network complexity and stability. Nature Climate Change, 11(4), 343–348. 10.1038/s41558-021-00989-9

Zhou, J., Xue, K., Xie, J., Deng, Y., Wu, L., Cheng, X., Fei, S., Deng, S., He, Z., Van Nostrand, J. D., & Luo, Y. (2012). Microbial mediation of carbon-cycle feedbacks to climate warming. Nature Climate Change, 2(2), 106–110. 10.1038/nclimate1331

Zhou, Y., Sun, B., Xie, B., Feng, K., Zhang, Z., Zhang, Z., Li, S., Du, X., Zhang, Q., Gu, S., Song, W., Wang, L., Xia, J., Han, G., & Deng, Y. (2021). Warming reshaped the microbial hierarchical interactions. Global Change Biology, 27(24), 6331–6347. 10.1111/gcb.15891

Zhou, Y., Wang, H., Xu, S., Liu, K., Qi, H., Wang, M., Chen, X., Berg, G., Ma, Z., Cernava, T., & Chen, Y. (2022). Bacterial-fungal interactions under agricultural settings: From physical to chemical interactions. Stress Biology, 2(1), 22. 10.1007/s44154-022-00046-1

Zosso, C. U., Ofiti, N. O. E., Soong, J. L., Solly, E. F., Torn, M. S., Huguet, A., Wiesenberg, G. L. B., & Schmidt, M. W. I. (2021). Whole-soil warming decreases abundance and modifies the community structure of microorganisms in the subsoil but not in surface soil. SOIL, 7(2), 477–494. 10.5194/soil-7-477-2021

